# Disassembly and degradation of MinD oscillator complexes by *Escherichia coli* ClpXP

**DOI:** 10.1101/2020.01.09.899195

**Authors:** Christopher J. LaBreck, Catherine E. Trebino, Colby N. Ferreira, Josiah J. Morrison, Eric C. DiBiasio, Joseph Conti, Jodi L. Camberg

## Abstract

MinD is a cell division ATPase in *Escherichia coli* that oscillates from pole to pole and regulates the spatial position of the cell division machinery. Together with MinC and MinE, the Min system restricts assembly of the FtsZ-ring to midcell, oscillating between the opposite ends of the cell and preventing FtsZ-ring misassembly at the poles. Here, we show that the ATP-dependent bacterial proteasome complex ClpXP degrades MinD in reconstituted degradation reactions in vitro, through direct recognition of the MinD N-terminal region, and in vivo. MinD degradation is enhanced during stationary phase, suggesting that ClpXP regulates levels of MinD in cells that are not actively dividing. MinC and MinD are known to co-assemble into linear polymers, therefore we monitored copolymers assembled in vitro after incubation with ClpXP and observed that ClpXP promotes rapid MinCD copolymer disassembly as a result of direct MinD degradation by ClpXP. The N-terminus of MinD, including residue Arg 3, which is near the ATP-binding site, is critical for degradation by ClpXP. Together, these results demonstrate that ClpXP degradation modifies conformational assemblies of MinD in vitro and depresses Min function in vivo during periods of reduced proliferation.

## Introduction

Cytokinesis in prokaryotes is a highly organized cellular process wherein a network of widely conserved cell division proteins function together to divide a single bacterial cell into two identical daughter cells (1). In *E. coli*, cell division commences with the assembly of a large ring-like protein structure termed the Z-ring, which contains bundled polymers of the GTPase FtsZ, and FtsZ-interacting proteins including FtsA and ZipA, and serves as the site of constriction (2). The Z-ring is a highly dynamic structure wherein FtsZ subunits are rapidly exchanged with a cytoplasmic pool via cycles of GTP binding and hydrolysis (3). Many proteins interact with FtsZ to spatially and temporally regulate Z-ring assembly, and a number of these proteins modulate FtsZ dynamics in the Z-ring (4).

The Min system of *Escherichia coli* functions to spatially regulate the site of cell division by inhibiting establishment of the Z-ring near the cell poles. MinD is one of three components of the Min system of proteins in *E. coli*, which includes MinC, MinD and MinE. These proteins oscillate across the longitudinal axis of the cell to prevent Z-ring assembly at the poles in *E. coli* (5). The Min system is used by several taxa to regulate division-site selection; however, the oscillation observed in *E. coli* is not preserved across all organisms that contain a Min system, and some organisms lack a Min system entirely (5). MinC directly interacts with FtsZ to disrupt GTP-dependent polymerization in vitro (6,7). The cellular distribution of MinC is determined by MinD via a direct protein-protein interaction. MinD is a member of the Walker A cytoskeletal ATPases (WACA) protein family and contains a deviant Walker A motif (8,9).MinD associates with the cytoplasmic membrane in an ATP-bound dimer conformation via a C-terminal membrane targeting sequence (MTS). MinE binds to MinD, stimulating ATP hydrolysis and displacement of MinD from the membrane (10,11).

MinC and MinD from several organisms, including *E. coli*, assemble into ATP-dependent cofilaments in vitro (12–14). The Lowe group solved a crystal structure of the *Aquifex aeolicus* MinCD complex, which supports a model in which *A. aeolicus* copolymers contain alternating MinC and MinD dimers (12,13). In *E. coli*, residues on the surface of the MinC C-domain (CTD) are important for copolymerization with MinD (7,12). Although several groups have reported copolymer formation in vitro, the physiological consequences of MinCD assembly in vivo are largely unknown. The Lutkenhaus group reported that MinD mutants that fail to polymerize with MinC, but still interact with MinC at the membrane, do not result in functional defects in vivo (15). Although copolymers are not essential to complete division in vivo, their assembly likely modifies Min patterning in vivo through direct competition of accessible MinD surfaces by MinC and MinE (7).

In several prokaryotes, cytokinesis is regulated proteolytically by the two-component ATP-dependent protease ClpXP (16). In *E. coli*, targeted degradation of FtsZ by ClpXP modulates Z-ring dynamics during the division process (4,17,18). Additional *E. coli* cell division proteins have also been identified as ClpXP proteolysis substrates, including ZapC (19) and MinD, which was previously implicated as a substrate in a proteomics study performed under DNA damage conditions (20,21). ClpXP contains both an unfoldase, ClpX, which is a hexameric ring-like AAA+ (ATPase Associated with diverse cellular Activities) ATPase, and the compartmentalized serine protease, ClpP, which is composed of two stacked heptameric rings (22). The ClpX unfoldase contains an N-terminal substrate binding domain, also called the zinc-binding domain (ZBD), that can undergo dimerization in solution and engages some substrates, including FtsZ and phage lambda O protein (λO), in addition to substrate specific adaptor proteins, such as SspB (18,23–25). After engagement of a substrate, ClpX utilizes ATP hydrolysis to unfold and translocate substrates through its axial channel into the central chamber of ClpP for degradation (26,27). Here, to determine if *E. coli* ClpXP regulates Min system function in *E. coli* by direct degradation of MinD and understand how ClpX targets MinD, we reconstituted in vitro degradation assays with MinD and MinD-containing complexes, including MinCD copolymers. We show that ClpXP degrades MinD and disassembles MinCD copolymers in vitro. We further demonstrate that ClpXP modifies Min function and oscillation in vivo.

## Results

### ClpXP degrades MinD in vitro

MinD was previously identified as a substrate for ClpXP degradation in *E. coli* (20). In addition, deletion of either *clpX* or *clpP* from a *minC* deletion strain leads to a synthetic filamentous phenotype during exponential growth (28). FtsZ, the major component of the Z-ring, is also degraded by ClpXP and an imbalance of FtsZ levels leads to filamentation (4,18). To further understand how degradation of MinD by ClpXP impacts Min function during division, we first developed an in vitro degradation assay using purified proteins. MinD (6 μM) was incubated with ClpX (1.0 μM), ClpP (1.2 μM) and ATP for 3 h, and degradation was measured by monitoring the loss of full-length MinD in the reaction with time (Fig. 1A). After incubation of MinD with ClpXP and ATP, we detected 45.9% less MinD after 3 hours (Fig. 1A); however, when either ATP or ClpP was omitted from reactions, the level of MinD did not change over the course of the experiment indicating that MinD is degraded by ClpXP in an ATP-dependent manner (Fig. 1A).

**Fig. 1.**
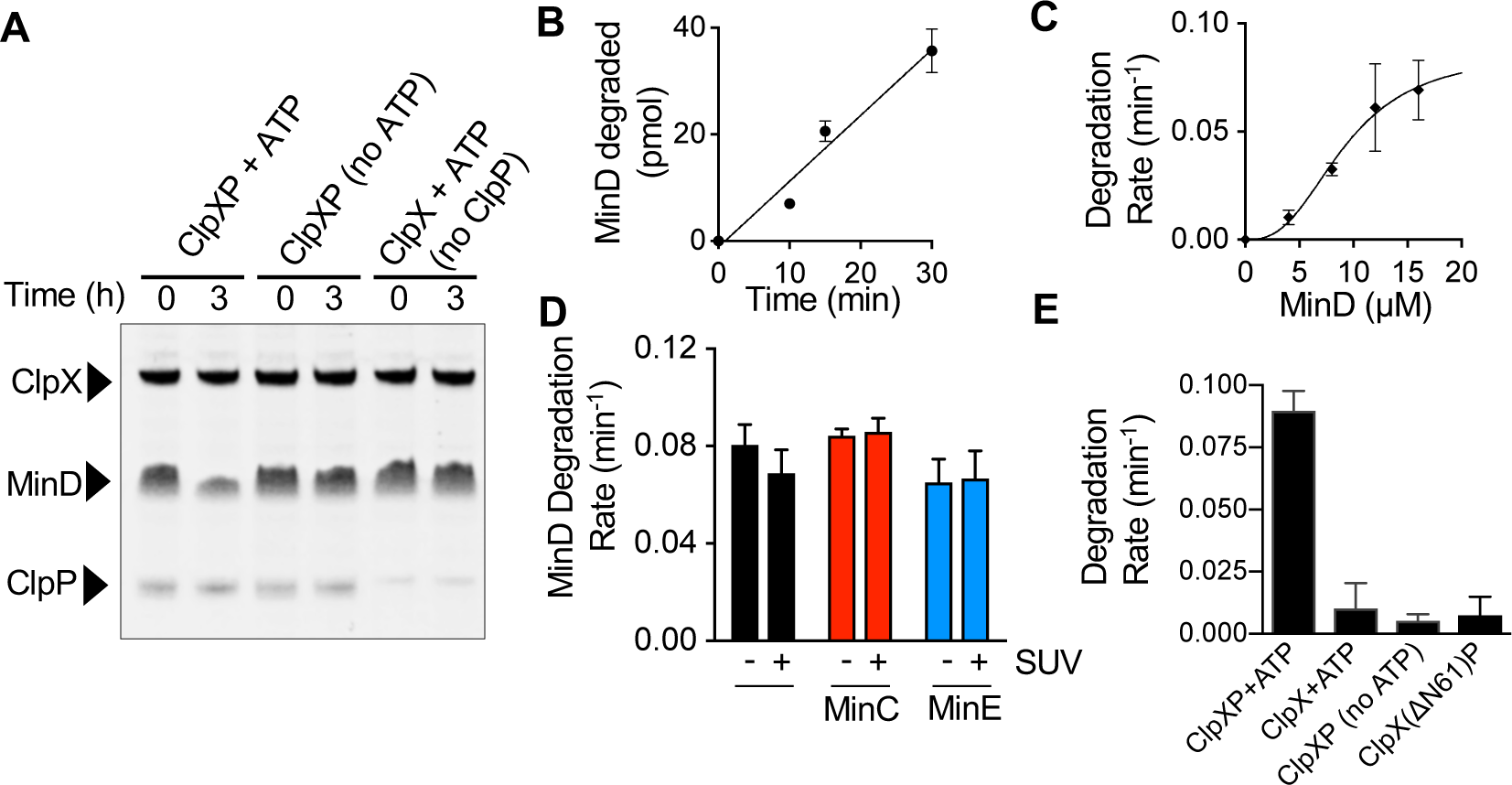
ClpXP degrades MinD in vitro. (A) MinD degradation was measured by monitoring the loss of MinD protein after 3 h in reactions containing MinD (6 μM), ClpX (1.0 μM), ClpP (1.2 μM), ATP (8 mM) and an ATP regenerating system, where indicated. (B) Degradation of fluorescent MinD was monitored in reactions containing ClpXP (0.8 μM), MinD (10 μM) and ATP (8 mM) with a regenerating system for 30 min. Degradation products were collected by ultrafiltration and quantified by fluorescence. (C) The rate of MinD degradation was determined for reactions containing ClpXP (0.8 μM), MinD (0 to 16 μM) and ATP (8 mM) and a regenerating system. (D) MinD degradation was monitored in reactions containing ClpXP (0.8 μM), ATP (8 mM) and a regenerating system, with MinD (10 μM), MinC (5 μM), MinE (10 μM), and SUVs (0.25 mg ml^−1^), where indicated. (E) The rate of MinD degradation by ClpXP or ClpX(ΔN61) with ClpP was determined for reactions containing MinD (10 μM), ClpP (0.9 μM), ATP (8 mM), and ClpX (0.75 μM) or ClpX (ΔN61) (0.75 μM), where indicated. Data from at least 3 replicates are shown as mean ± SEM.

To quantitatively measure the rate of degradation, we labeled MinD with Alexa fluor 488 and measured degradation by monitoring fluorescent peptides released following incubation with ClpXP and ATP. Degradation reactions containing ClpXP (0.7 μM), ATP (8 mM), and MinD (10 μM) were stopped by the addition of EDTA (50 mM), and fluorescent peptides were collected by ultrafiltration and quantified by fluorescence. We observed that the amount of MinD degraded increased linearly over the course of 30 min (Fig. 1B). Next, we examined the rate of MinD degradation by ClpXP with increasing MinD concentration (0 to 16 μM). We observed a concentration-dependent increase in the rate of MinD degradation, which plateaus near 20 μM MinD, with a rate of 0.08 ± .01 min^−1^ (Fig. 1C). The degradation rate of another substrate, FtsZ, has also been shown to increase with increasing substrate concentration, which suggests a low affinity interaction at low substrate concentrations (4,18). Together, these results demonstrate that ClpXP recognizes and degrades MinD in a concentration-dependent manner.

In the presence of ATP, MinD binds to *E. coli* phospholipids by inserting a C-terminal amphipathic helix into the phospholipid bilayer and recruits MinC to phospholipids in vitro. MinE also binds to membrane-associated MinD in vitro. MinE stimulates ATP hydrolysis by MinD in the presence of phospholipids, and then MinD dissociates from the phospholipid bilayer (29,30). Since MinD is capable of binding directly to MinC, MinE, and phospholipids, we tested if the addition of *E. coli* phospholipids, prepared as small unilamellar vesicles (SUVs), modifies the rate of MinD degradation in absence and presence of MinC and MinE. We observed that the interaction with SUVs, MinC (10 μM) or MinE (10 μM) had no significant impact on the rate of MinD degradation by ClpXP under the conditions tested (Fig. 1D). Lastly, to confirm that ClpXP does not also degrade MinC or MinE, we monitored degradation of both MinC and MinE, but detected no proteolysis of either protein after 3 hours (Fig. S1).

The ClpX N-terminal domain is a zinc-binding domain (ZBD) that can dimerize independently of an attached ClpX AAA+ ATPase domain (25). ClpX(ΔN61) is an engineered deletion mutant protein that lacks the N-terminal 61 amino acids, which includes the ZBD, but retains the ability to form a functional complex with ClpP and degrade some substrates, including ssrA-tagged proteins independently of the SspB adaptor protein (31,32). Other substrates, including MuA tetramers, λO and FtsZ are targeted by the ClpX N-domain for recognition and degradation (18,25,31). To determine if the N-terminal domain of ClpX is important for recognition and degradation of MinD, we purified ClpX(ΔN61) and compared the rate of MinD degradation by ClpX(ΔN61)/ClpP to the rate of degradation by ClpXP. In reactions containing 10 μM MinD, we observed no detectable MinD degradation by ClpX(ΔN61)/ClpP, in contrast to ClpXP (Fig. 1E). Furthermore, both purified ClpX and ClpX(ΔN61) supported degradation of Gfp-ssrA with ClpP in vitro (Fig. S2A). Consistent with the requirement for the ZBD, we also found that a peptide, called XB, which contains the ZBD-interacting region of the SspB adaptor protein, competitively inhibits MinD degradation (Fig. S2B), suggesting that the MinD and SspB binding sites may overlap on the ClpX N-domain. Together, our results suggest that the ClpX N-domain is important for recognition and degradation of MinD.

### ClpXP antagonizes MinCD copolymers in vitro

MinC and MinD from *E. coli, Pseudomonas aeruginosa* and *Aquifex aeolicus*, readily form copolymers in the presence of ATP (12–14) (Fig. 2A). ClpXP is known to disassemble FtsZ polymers in vitro (4,18), therefore we tested if ClpXP could also prevent assembly or destabilize MinCD copolymers. First, to test if the presence of ClpXP in an assembly reaction reduces or prevents copolymerization of MinD with MinC in vitro, we monitored 90° light scatter of reactions containing MinD (8 μM), MinC (4 μM), and then added ATP alone or with ClpX and/or ClpP, where indicated (Fig. 2B and 2C). Light scatter was then monitored for an additional 30 min. The addition of ATP without ClpXP stimulated robust copolymer formation; however, the addition of ATP and ClpXP lead to a small increase in light scatter that rapidly decreased with time (Fig. 2B). Next, we tested if ClpX inhibits MinCD assembly without ClpP, since inhibition of MinCD copolymer assembly could potentially result from MinD unfolding, but not degradation. Interestingly, we found that the addition of ClpX resulted in a 45% inhibition of copolymerization (Fig. 2C), and also that the ClpX ATPase mutant protein, ClpX(E185Q), which hexamerizes and binds substrates but does not unfold them, similarly impaired copolymer formation (45% inhibition). Together, these results suggest that ClpX is capable of impairing MinCD assembly independently of ClpP and ATP hydrolysis, but that ClpXP is substantially more effective for preventing assembly (Fig. 2B and 2C). Addition of an equivalent volume of buffer alone does not inhibit copolymer formation in control experiments (Fig. 2B and 2C). Lastly, we also observed that ClpP without ClpX alone does not modify light scatter of MinCD copolymers with ATP (4 mM). Together, our results suggest that MinCD copolymer formation is reduced by ClpX and prevented by ClpXP.

**Fig. 2.**
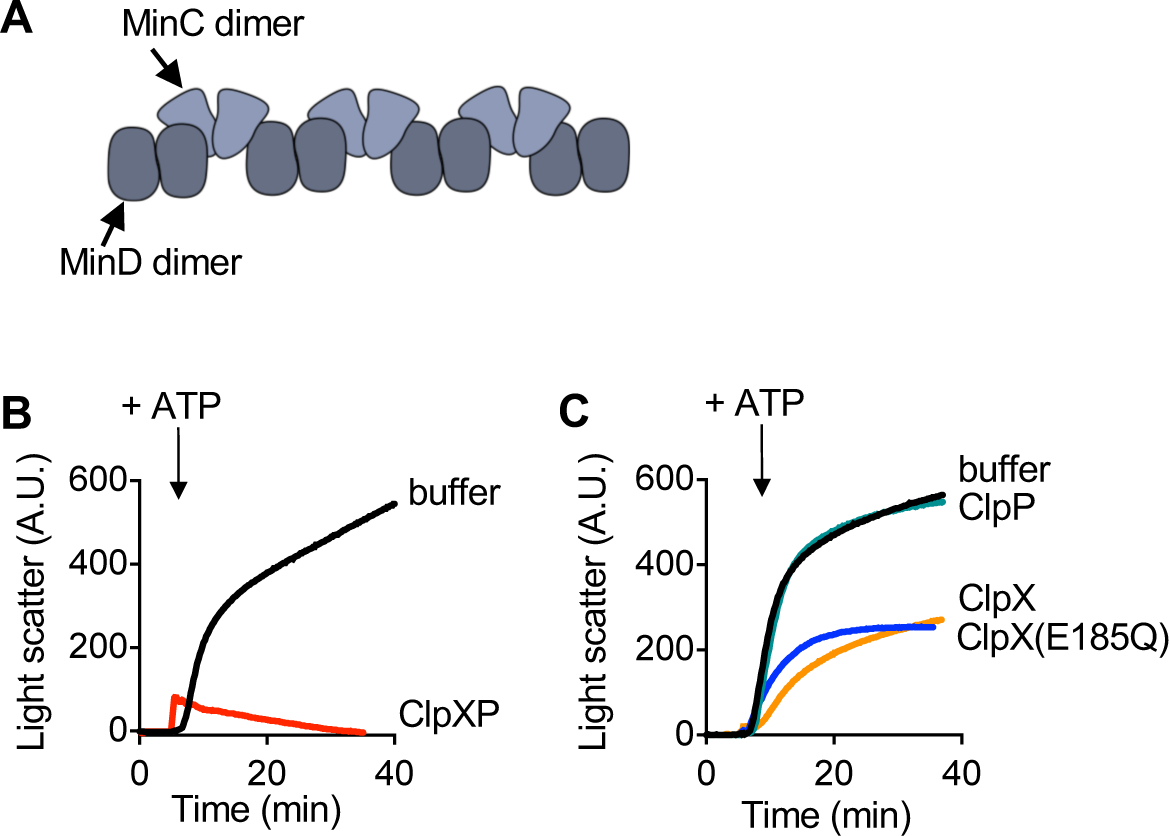
ClpXP prevents MinCD copolymer assembly. (A) MinC and MinD dimers assemble into alternating copolymers. (B) MinCD copolymer assembly was monitored by 90° light scatter in reactions containing MinC (4 μM), MinD (8 μM), ClpX (0.75 μM), and ClpP (0.9 μM), where indicated. After 5 min, ATP (8 mM) was added and reactions were monitored for an additional 30 min. (C) MinCD copolymer assembly was monitored by light scatter as described in (B) in reactions containing MinC (4 μM), MinD (8 μM), with either buffer, ClpX (0.75 μM), ClpX(E185Q) (0.75 μM) or ClpP (0.9 μM), where indicated. Data shown is representative of at least three replicates.

Next to determine if ClpXP destabilizes pre-assembled MinCD copolymers, we performed an order of addition experiment. First, copolymers were pre-assembled with ATP, and then ClpXP (0.5 to 0.9 μM) was added and copolymer disassembly was followed by light scatter for another 30 min. Addition of ClpXP led to a rapid decrease in light scatter that correlated with ClpXP concentration (Fig. 3A). In contrast, the addition of buffer, ClpX with ATP, or ClpP failed to promote disassembly of MinCD copolymers (Fig. 3B) (Fig. S3). Finally, we directly visualized copolymers via negative staining transmission electron microscopy (TEM) and compared MinCD copolymer abundance and morphology to copolymers incubated with ClpXP. Consistent with previous reports, we observed MinCD copolymers with ATP (4 mM) (Fig. 3C). Next, ClpXP (0.9 μM) with ATP (4 mM) alone and added to reactions containing MinCD copolymers pre-assembled with ATP were assembled. After 15 min, reaction products were analyzed by TEM. In the presence of ClpXP (Fig. 3C), we observed copolymers that were shorter and spread more sparsely across the grid (Fig. 3C), compared to copolymers without ClpXP. To confirm that polymers were not observed in reactions containing ClpXP alone, we visualized ClpXP (0.9 μM) assembled with ATP (4 mM) and a regenerating system. We detected a homogenous population of ClpXP particles (Fig. 3C) and did not observe any polymeric structures. In the structural model of MinCD, copolymers are comprised of alternating MinC and MinD dimers (12). Therefore, it is possible that copolymer disassembly by ClpXP could arise from degradation of MinC or a failure of MinC to dimerize and/or interact with MinD. However, we observed no degradation of MinC by SDS-PAGE (Fig. S1), therefore copolymer disassembly is likely not mediated by MinC degradation.

**Fig. 3.**
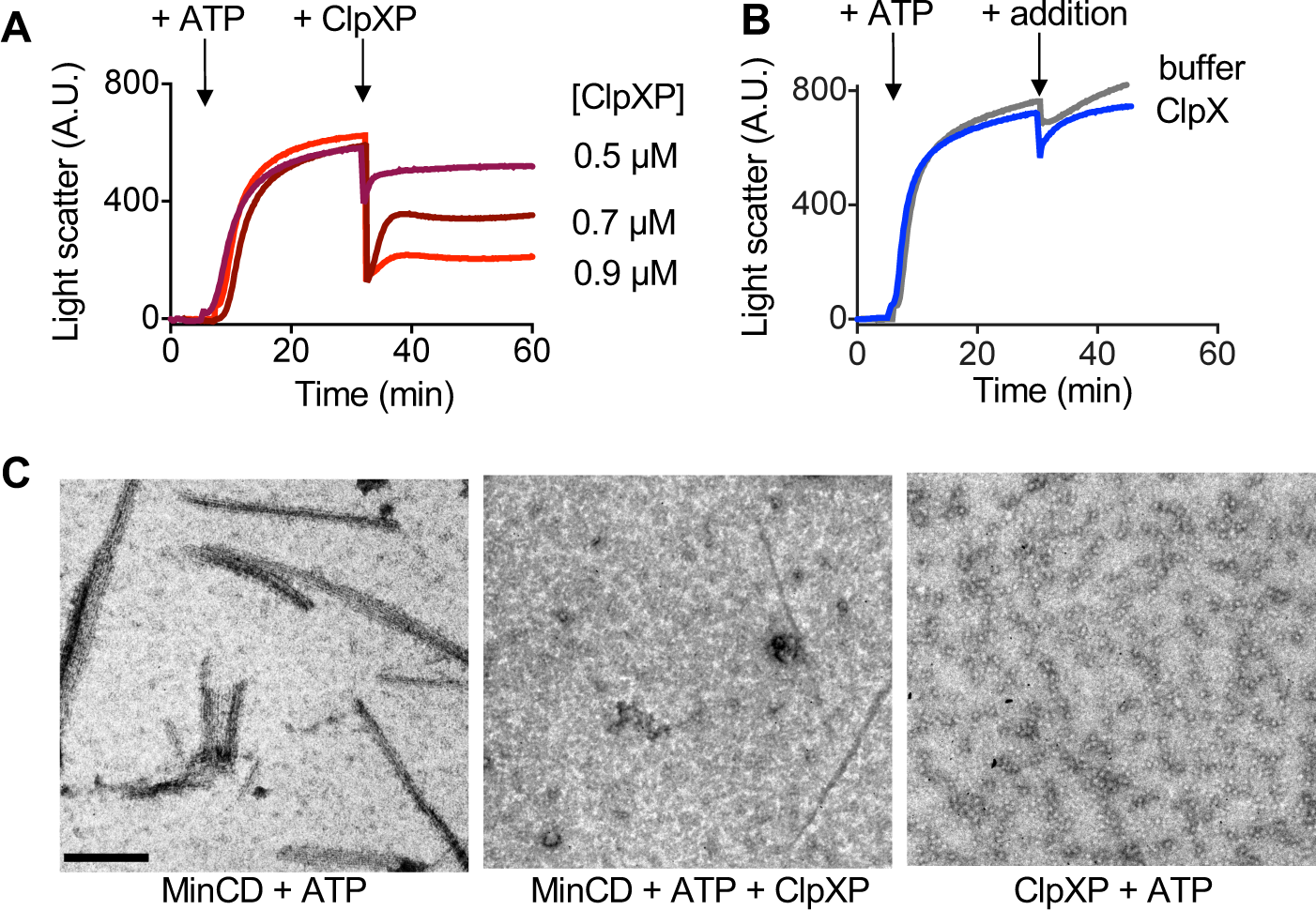
Monitoring disassembly of MinCD copolymers by ClpXP. (A) MinCD copolymers were assembled with MinC (4 μM), MinD (8 μM), and ATP (8 mM) and monitored by 90° light scatter. After 30 min, where indicated, ClpXP (0.5 to 0.9 μM), (B) ClpX (0.75 μM) alone or buffer was added. Light scatter was monitored for an additional 30 min. In (A) and (B) curves are representative of at least three replicates. (C) Polymers were visualized by TEM in reactions containing MinC (4 μM), MinD (8 μM), and ATP (8 mM) with and without ClpXP (0.75 μM) and ClpP (0.9 μM), where indicated. Reactions were applied to carbon-coated grids, fixed with glutaraldehyde and stained with uranyl-acetate. Scale bar is 200 nm.

### The MinD N-terminal sequence contains residues that are important for degradation by ClpXP

Substrate recognition by ClpX is mediated by the presence of different sequence motifs, or degrons, at the N-or C-terminal regions of protein substrates (21,33). The N-terminus of MinD contains amino acids similar to the N motif-2 consensus motif (M-b-ϕ-ϕ-ϕ- X5-ϕ) for ClpX (20,21) (^1^MARIIV-X_5_-G^12^). In the structural model of MinD, Arg 3, is present near the N-terminus and accessible to the surface (Fig. 4A). To determine if this arginine is important for recognition by ClpX, we mutagenized the residue to glutamate and purified MinD(R3E). In degradation reactions with ClpXP in vitro, we observed that 50% of wild type MinD was degraded in the first 60 min; however, we detected no MinD(R3E) degradation during the experiment under the conditions tested (Fig. 4B). To confirm that MinD(R3E) is not defective for function, we measured the ability of MinE to stimulate ATP hydrolysis of MinD(R3E) in the presence of SUVs. We observed that the ATP hydrolysis rate of MinD(R3E) (8 μM) was stimulated 10-fold by MinE (16 μM) and SUVs (1 mg ml^−1^), similar to wild type MinD, suggesting that the amino acid substitution does not impair MinD function (Fig. 4C).

**Fig. 4.**
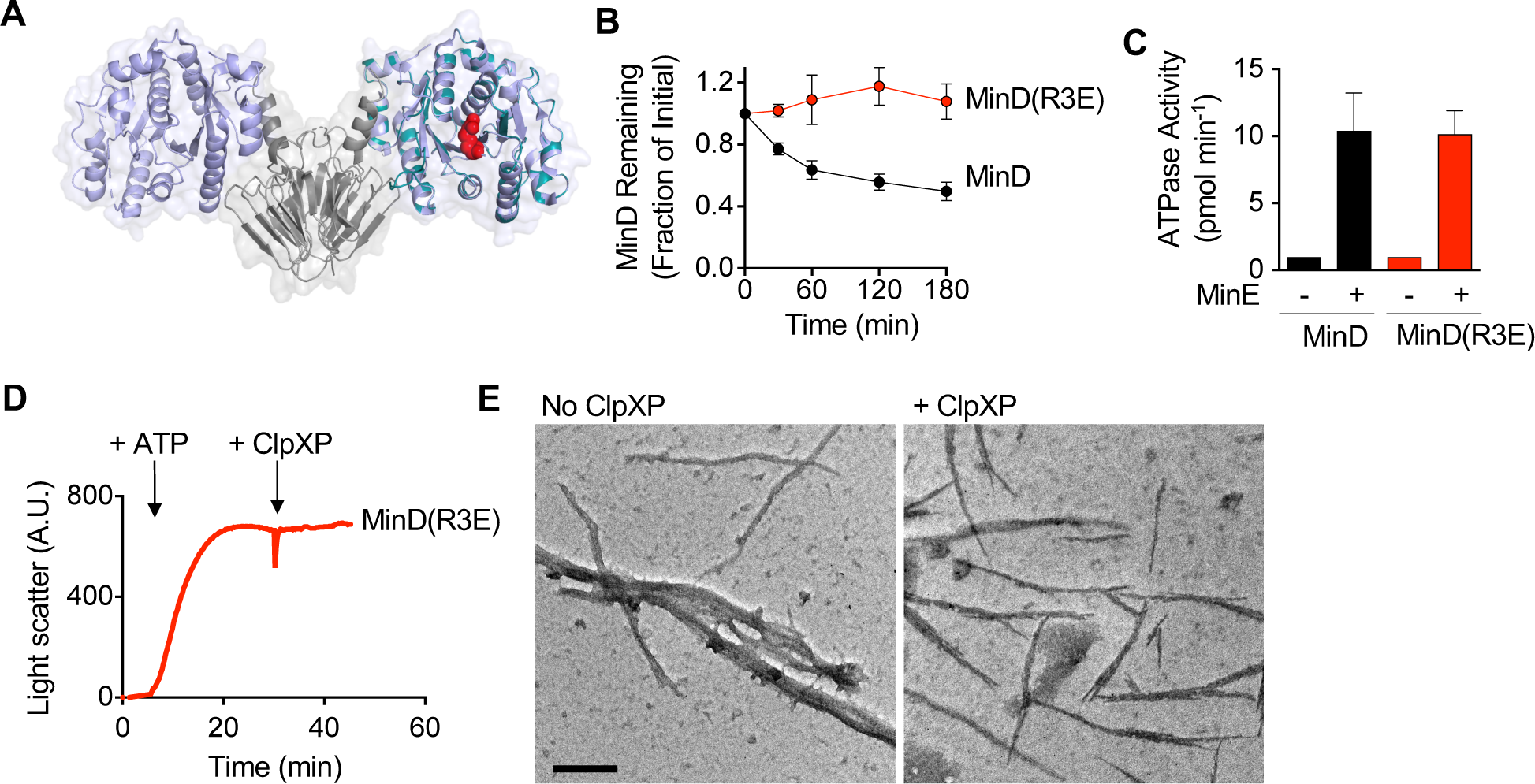
MinD(R3E) is defective for degradation by ClpXP. (A) Arg 3 of MinD is located near the ATP-binding site of MinD. *E. coli* MinD (cyan) was modeled onto *Aquifex aeolicus* MinD (purple) in complex with MinC C-domain (gray) (pdb 4V02) (12). (B) Degradation was monitored in reactions containing ClpX (0.75 μM), ClpP (0.9 μM), ATP (8 mM), an ATP regenerating system, and MinD (8 μM) or MinD(R3E) (8 μM), where indicated. (C) ATP hydrolysis by MinD(R3E) with and without stimulation by MinE and SUVs was measured by monitoring phosphate release over 10 min in reactions containing SUVs (1 mg ml^−1^), ATP (4 mM), and MinD (8 μM), MinD(R3E) (8 μM), and MinE (16 μM), where indicated. (D) Copolymer formation by MinD(R3E) with MinC was compared by monitoring 90° light scatter with ATP for 30 min, and then ClpXP was added to the reaction and light scatter was monitored for an additional 30 min. (E) Polymers containing MinC and MinD(R3E) were visualized by TEM in reactions containing MinC (4 μM), MinD(R3E) (8 μM), and ATP (8 mM) with and without ClpXP (0.75 μM) and ClpP (0.9 μM), where indicated. Scale bar is 200 nm.

MinD(R3E) is defective for degradation by ClpXP, but copolymerizes with MinC (Fig. S4A), therefore, we tested if whether copolymers containing MinD(R3E) and MinC are resistant to disassembly by ClpXP. As expected, we observed that ClpXP failed to destabilize pre-assembled MinC/MinD(R3E) polymers, in contrast to copolymers formed with MinC (4 μM) and wild type MinD (8 μM) (Fig. 4D, Fig. 3A and S4A). Our results suggest that ClpXP promotes disassembly of MinCD copolymers and that Arg 3 of MinD is important for ClpXP-dependent disassembly. Furthermore, MinD(R3E) (4 μM) recruited similar levels of MinC (4 μM) to SUVs compared to MinD (4 μM) (Fig. S4B) suggesting that it is not defective for known interactions in vitro. Finally, we visualized MinD(R3E)-dependent copolymers by TEM that were incubated with and without ClpXP (0.9 μM). As expected, copolymers were detected under both conditions, consistent with a failure of ClpXP to disassemble copolymers containing MinD(R3E) and MinC (Fig. 4E).

### MinD degradation and function in *clp* deletion strains

To monitor protein turnover of MinD in the K-12 strain MG1655 and to determine if ClpXP actively degrades MinD in vivo as well as in vitro, we performed an antibiotic chase experiment using spectinomycin to halt new protein synthesis and then monitored MinD levels by immunoblot. We observed that MinD is degraded in MG1655 cells from 16 h cultures (OD_600_ of 2.0 A.U.) and with a long half-life of approximately 180 min (Fig. 5A) (Fig. S5A). However, deletion of *clpX* or *clpP* prevents the degradation and significantly stabilizes the protein over the course of the experiment, indicating that ClpXP is predominantly responsible for the observed turnover in this strain under the conditions tested (Fig. 5A) (Fig. S5A). Surprisingly, in log phase cells (OD_600_ of 0.3 A.U.), we observed even slower MinD protein turnover relative to stationary phase, with MinD persisting much longer than 180 min and similar to a *clpX* deletion strain (Fig S5B).

**Fig. 5.**
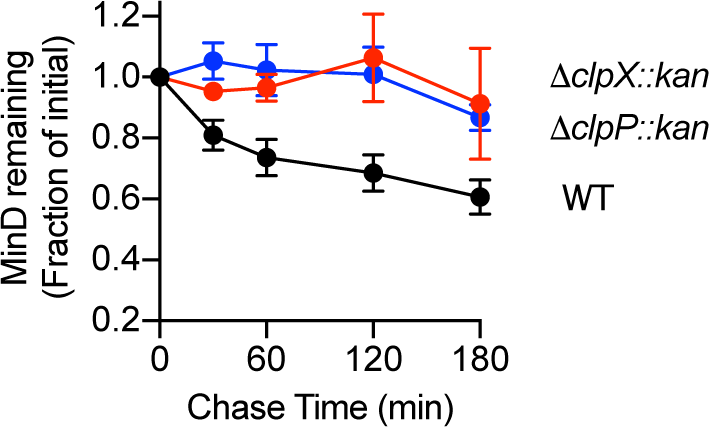
MinD degradation in vivo by ClpXP. Relative MinD levels were monitored in cells that were grown to stationary phase in LB and then treated with spectinomycin (200 μg ml^−1^). Cell extracts from wild type MG1655, *clpX* and *clpP* deletion strains were analyzed by immunoblotting for MinD followed by densitometry.

In live cells, MinD levels during log phase or stationary phase are controlled by a balance of synthesis and degradation, with partitioning of Min proteins into daughter cells during division events. Therefore, we next tested if Min oscillations are more active or detectable in log phase versus stationary phase. To monitor MinD oscillation in vivo, we replaced *minD* in the chromosomal *min* operon with *gfp-minD* (Table 1). This gene replacement resulted in cells with a mean cell length of 1.92 ± 0.04 μm (n = 200), which is similar to wild type cells with a mean cell length of 1.93 ± 0.03 μm (n = 200), suggesting that Gfp-MinD is fully functional for regulating division. We monitored productive oscillations in cells expressing Gfp-MinD and found that over half of the cells (51%) harvested from log phase cultures (OD_600_ of 0.3 A.U.) showed productive MinD oscillations. In contrast, only 38% of cells expressing Gfp-MinD harvested from stationary phase 16 h cultures (OD_600_ of 2.0 A.U.) showed productive oscillations, suggesting that the Min system is depressed in cells that are in stationary phase. Since MinD degradation occurs during stationary phase, these results suggest that ClpXP may contribute to reducing MinD levels and function in slow-growing cells, leading to fewer cells that exhibit Min oscillation.

**Table 1.** *E. coli* strains and plasmids.

| Strain or Plasmid | Genotype | Source, reference or construction |
| --- | --- | --- |
| **Strains** |  |  |
| MG1655 | *LAM-rph-1* | (46) |
| BL21 (λDE3) | *F– ompT gal dcm lon hsdSB(rB-mB-) λ(DE3[lacI lacUV5-T7 gene 1 ind1 sam7 nin5])* | EMD Millipore, USA |
| CL0030 | MG1655 Δ*minC::kan-Prha-parE* | (7) |
| CL0428 | MG1655 Δ*minC::Pbad-gfp-minC* | (7) |
| CF020 | MG1655 Δ*minC::Pbad-gfp-minC*<br>Δ*clpPX::kan-Prha-parE* | CL0428; λred |
| CL0476 | MG1655 Δ*minD::kan-Prha-parE* | MG1655; λred |
| CF005 | MG1655 Δ*minD::gfp-minD* | CL0476; λred |
| CF015 | MG1655 Δ*minD::gfp-minD*<br>Δ*clpPX::kan-Prha-parE* | CF005; λred |
| JC0259 | MG1655 Δ*clpX::kan* | (28) |
| JC0263 | MG1655 Δ*clpP::kan* | (28) |
| **Plasmids** |  |  |
| pEt-MinC | *kan* | (13) |
| pEt-MinD | *kan* | (13) |
| pKD46 | *amp*; λred recombinase | (42) |
| pKD267 | *kan-Prha-parE* | (7) |
| pBAD33-*gfp* | *cat* | (7) |
| pBAD33-*gfp-minC* | *cat* | (7) |
| pBAD33-*gfp-minD* | *cat* | This study |
| pBR322 | *amp* |  |
| pClpXP | *amp* | (18) |

Finally, to determine if ClpXP degradation activity modifies the Min localization pattern in cells during productive oscillations in log phase, we measured the rate of fluorescent foci movement from pole to pole by Gfp-MinD in cells with and without ClpXP. We observed polar oscillation of Gfp-MinD in live, dividing cells with movement of the fluorescent foci across the longitudinal axis of the cell at a rate of 160.4 ± 7.6 nm sec^−1^ (Fig. S6A and S6B). In cells deleted for *clpP* and *clpX* (Table 1), which is present in a single operon and denoted here as *clpPX*, we observed a modest, but significant 15% reduction in the Gfp-MinD oscillation rate compared to cells with *clpPX* intact (Fig. S6A and S6B). The rate of oscillation by Gfp-MinD is within error of the oscillation of Gfp-MinC, measured here at 163.3 ± 4.8 nm sec^−1^ (Fig. S6C), and reported previously (7). Deletion of *clpPX* also led to a 15% reduced Gfp-MinC oscillation rate, which is dependent on MinD for oscillation (Fig. S6C). Surprisingly, overexpression of ClpXP from a vector that was previously shown to increase ClpX and ClpP levels by 4- and 75-fold, respectively, increased the oscillation rates of both Gfp-MinD and Gfp-MinC (Fig. S6B and S6C). Together, these results indicate that ClpXP contributes to setting MinC and MinD oscillation rates in dividing cells in vivo, either directly by managing MinD levels or indirectly by degradation of FtsZ and subsequently impacting Min function.

## Discussion

Here, we demonstrate that the two component ATP-dependent protease ClpXP degrades the cell division protein MinD in vitro. Degradation of MinD requires the ClpX N-domain and utilizes a MinD N-terminal region that has a sequence similar to other substrates with predicted N-terminal degrons (Fig. 1). We identified a residue in the MinD N-terminal region, Arg 3, that is important for recognition and degradation by ClpXP. Although ClpX functions with ClpP to regulate overall protein turnover of many substrates, such as RpoS in vivo, ClpX also remodels protein substrates. For example, ClpX remodels MuA to regulate phage transposition and can function as a protein disaggregase (34–36). Given that ClpX is capable of remodeling protein complexes with and without ClpP, we investigated the ability of ClpX and ClpXP to modulate MinCD copolymer assembly in vitro. We found that ATP-driven coassembly of MinCD was partially prevented in the presence of ClpX and abolished in the presence of ClpXP (Fig. 2B). Furthermore, copolymer formation was similarly impaired by ClpX and ClpX(E185Q), which contains a mutation in the Walker B motif and is defective for ATP hydrolysis, but still oligomerizes and binds to substrates (37). Thus, ClpX alone antagonizes MinCD copolymer assembly via an ATP hydrolysis-independent mechanism. Consistent with these results, previous studies demonstrated that in *B. subtilis*, ClpX impairs FtsZ polymer assembly through an ATP-independent mechanism (38,39). ATP-hydrolysis by ClpX is required for substrate unfolding and translocation but not substrate binding. Therefore, our results are consistent with ClpX inhibiting copolymerization via a binding/holding mechanism, where the direct interaction, without unfolding, may be sufficient to impair assembly or “cap” copolymers.

In *E. coli*, ClpXP degrades FtsZ to modulate Z-ring subunit exchange and dynamics during division in vivo (4,18). Eukaryotic members of the AAA+ protein family, including spastin and katanin, are also capable of microtubule polymer disassembly (40). Thus, a major function of ClpXP may be to disassemble polymer networks in the cytoplasm. MinD and MinC from several organisms form large linear filaments that are readily observed by electron microscopy (7,12–14). *E. coli* MinCD copolymers are rapidly disassembled by ClpXP in vitro (Fig. 2 and 3). To date, multimerizing protein substrates, including FtsZ, MuA, DPS and MinD, rely at least partly on the ClpX N-domain for efficient remodeling and/or degradation, which implicates a role for the ClpX N-domain in coordinating multiple substrate interactions with substrate engagement and initiation of unfolding.

Gfp-MinD oscillation occurs in the absence of MinC, and cells expressing MinD mutant proteins that are impaired for copolymerization with MinC do not have obvious cell division defects in vivo (15,41). However, copolymerization with MinC may limit the available population of MinD that is activated in vivo through sequestration in the cytoplasm or concentration on the membrane thereby modifying oscillation. We observed that ClpXP, and to a lesser extent ClpX, directly alters copolymer assembly and abundance in vitro (Fig. 2 and 3). Thus, ClpXP may also modulate the accessible population of MinD by modifying copolymer conformation. Surprisingly, when we tracked total MinD protein turnover, we observed that MinD was relatively stable, especially during log phase, yet susceptible to ClpXP proteolysis during stationary phase. It is currently unclear if the stationary phase MinD proteolysis that we observed serves a general housekeeping function of ClpXP, or if ClpXP is disassembling and degrading MinD-containing complexes to affect divisome function or alleviate a burden of intracellular ATP utilization consumed by MinD oscillation. Moreover, there appears to be variability in the extent of MinD degradation by ClpXP among strains and growth conditions, and MinD degradation is enhanced under DNA damaging conditions (20). Alterations in ClpXP expression levels modify Min oscillation rates in dividing cells, but it is unknown if this is a direct effect (e.g. resulting from MinD degradation and/or MinD complex disassembly) or this is an indirect effect (e.g. resulting from modified FtsZ levels). It is currently clear that ClpXP degradation fine tunes division in *E. coli* via degradation of FtsZ, ZapC and MinD, yet there is much that remains to be understood about the spatiotemporal coordination of these events, bulk effects on FtsZ polymer mass and dynamics and effects on cell cycle timing, constriction or septation.

## Experimental procedures

### Strain construction

Strains and plasmids are described in Table 1. Strains deleted for *clpX, clpP*, or the *clpPX* operon, and strains expressing Gfp-MinD, Gfp-MinC and MinD(R3E) from the native loci in the chromosome were constructed using Lambda Red recombination and gene replacement, where indicated, using negative selection by induction of *parE* by rhamnose, as described (7,42).

### Protein expression and purification

MinC, MinD, MinE, FtsZ, ClpX, ClpX(ΔN61), and ClpP, were each overexpressed from a plasmid in *E. coli* BL21 (λDE3) and purified as native proteins according to previously published methods (7,13,18,43). MinD and ClpX mutant proteins were constructed by site-directed mutagenesis of plasmids containing *minD* or *clpX* using the QuickChange II XL Site-directed mutagenesis kit (Agilent) and confirmed by sequencing prior to purification. Gfp-ssrA was purified as previously described (44). Protein concentrations are reported as MinC dimers, MinD dimers, MinE dimers, ClpX hexamers and ClpP tetradecamers.

### Phospholipid recruitment assays with MinC and MinD

MinD (4 μM) or MinD(R3E) (4 μM), where indicated, were added to reactions containing SUVs (0.25 mg ml^−1^), MinC (4 μM), and ATP (4 mM) in assembly buffer containing 50 mM 2-(*N*-morpholino)ethanesulfonic acid (MES)-pH 6.5, 100 mM KCl, 10 mM MgCl_2_. Small unilamellar vesicles from *E. coli* extracts (Avanti Polar lipids) were prepared as described (45). Following the addition of ATP, reactions were incubated for 10 min at 30 °C, and SUV-containing protein complexes were collected by centrifugation at 14,000 x *g* at 23 °C, resuspended in LDS loading buffer, and analyzed by SDS-PAGE and densitometry.

### ATP hydrolysis assays with MinD

ATP hydrolysis was measured by monitoring the amount of inorganic phosphate released after 10 min in reactions containing Tris buffer (20 mM, pH 7.5), KCl (50 mM), MgCl_2_ (10 mM), ATP (4 mM), MinD (8 μM), MinE (16 μM), and SUVs (1 mg ml^−1^), where indicated. Phosphate released in reactions was detected using Biomol green (Enzo Life Sciences) and compared to a phosphate standard curve.

### Degradation assays

FL-MinD (10 μM), or FL-MinD (4 to 16 μM), where indicated, was added to reactions containing ClpX (0.7 μM), ClpX(ΔN61) (0.7 μM), ClpP (0.8 μM), MinC (5 μM), MinE (10 μM), and ATP (10 mM) with an ATP regenerating system containing creatine phosphokinase (50 μg ml^−1^) and creatine phosphate (30 mM), where indicated, in degradation buffer (50 mM MES 6.5, 100 mM KCL, 10 mM MgCl_2_, 2 mM TCEP). Degradation reactions were incubated for 30 min at 30 °C and reactions were terminated by the addition of EDTA (50 mM). Degradation products were applied to pre-washed 3 kDa Nanosep spin-filters with Omega membrane (Pall). Samples were centrifuged at 14,000 x *g* for 20 min at 23 °C, and the fluorescence of the filtrate was measured with excitation and emission wavelengths set for 490 nm and 525 nm, respectively. Fluorescent MinD was generated by labeling with Alexa Fluor-488 succinimidyl ester (Life Technologies). Labeled MinD was tested for the ability to form ATP-dependent copolymers with MinC.

Degradation reactions using non-labeled MinD was monitored in reactions containing MinD (6 μM), MinD(R3E) (6 μM) or MinD(K11A) (6 μM), where indicated, MinC (6 μM), MinE (6 μM), where indicated, ClpX (1.0 μM), ClpP (1.2 μM), ATP (4 mM) and an ATP regenerating system containing creatine phosphokinase (50 μg ml^−1^) and creatine phosphate (30 mM), where indicated, in degradation buffer. Immediately following the addition of ATP, and after 30 min, 60 min, 120 min, and 180 min, or after 180 min, where indicated, samples were removed from degradation reactions and added to LDS loading buffer. Protein amounts were analyzed by SDS-PAGE and densitometry.

Degradation of Gfp-ssrA was monitored in reactions containing Gfp-ssrA (1 μM), ClpX (0.3 μM) or ClpX(ΔN61) (0.3 μM), ClpP (0.4 μM) ATP (4 mM) and an ATP regenerating system containing creatine phosphokinase (25 μg ml^−1^) and creatine phosphate (15 mM) in degradation buffer. Fluorescence was monitored with time using excitation and emission wavelengths set to 395 nm and 509 nm, respectively.

### Copolymer assays with MinC and MinD

To monitor MinCD copolymer formation by light scatter, MinD (8 μM) was added to reactions containing MinC (4 μM) in copolymer assembly buffer (20 mM MES, pH 6.5, 100 mM KCl, 10 mM MgCl_2_). 90° light scatter was monitored with excitation and emission wavelengths set to 450 nm. After 5 min, ATP (8 mM) or ClpX (0.75 μM), ClpX(E185Q) (0.75 μM), ClpP (0.9 μM), or ClpX (0.75 μM) and ClpP (0.9 μM), where indicated, and ATP (8 mM) and ATP regenerating system containing creatine phosphokinase (50 μg ml^−1^) and creatine phosphate (30 mM), were added and 90° light scatter was monitored for an additional 30 min. In reactions monitoring copolymer disassembly, 90° light scatter was initially monitored in reactions containing MinD (8 μM) or MinD(R3E) (8 μM) where indicated, MinC (4 μM), and ATP (4 mM) for 30 min at 23 °C. Then, ClpX (0.75 μM), ClpP (0.9 μM) or ClpX (0.4 to 0.75 μM) or ClpP (0.5 to 0.9 μM), where indicated, ATP (4 mM) and ATP regenerating system containing creatine phosphokinase (50 μg ml^−1^) and creatine phosphate (30 mM) were added and 90° light scatter was monitored for an additional 15 min.

### Electron microscopy

Reactions containing MinC (4 μM), MinD (8 μM) and MinD(R3E) (8 μM), where indicated, ClpX (0.75 μM) and ClpP (0.90 μM), where indicated, ATP (8 mM) and ATP regenerating system containing creatine phosphokinase (50 μg ml^−1^) and creatine phosphate (30 mM) in copolymer assembly buffer were applied to 300-mesh carbon/formvar-coated grids, fixed with glutaraldehyde (2%) and stained with uranyl acetate (2%). Samples were imaged by transmission electron microscopy using a JEM-2100 80 Kev instrument.

### Fluorescence microscopy

Wild-type, overexpression, and deletion strains of *gfp-minC* and *gfp-minD* were grown, and oscillation rates observed and plotted as described previously (7). Strains containing pBR322 and pClpXP were grown in the presence of ampicillin in order to maintain the presence of the plasmids.

### Antibiotic chase and immunoblotting

Stationary phase bacterial cultures in Luria-Bertani (LB) medium at OD_600_ 0.05 where grown at 30 °C for 16 h. Log phase cultures were grown overnight and diluted into fresh LB medium OD_600_ 0.05 and grown to OD_600_ of 0.3 A.U. at 30 °C. One ml or 5 ml samples respectively were collected at 0, 30, 60, 120, and 180 min. Spectinomycin (Sigma) (200 μg ml^−1^) was added at 0 min. Proteins were precipitated with 15% trichloroacetic acid (Sigma) for 30 minutes at 4 °C. Suspensions were then centrifuged at 5,000 x *g* for 10 min at 4^0^C. Pellets were isolated and washed with acetone for 10 min at 4^0^C followed by centrifugation at 10,000 x *g* for 10 min at 4 °C. Acetone was removed and pellets were resuspended in 10% sodium dodecyl sulfate (SDS). Protein samples were analyzed by reducing SDS-PAGE and transferred to a nitrocellulose membrane (Invitrogen). Membranes were washed with tris buffered saline (pH 7.6) and Tween-20 (0.05%) (TBST), blocked for 2 hours with 2% (w/v) bovine serum albumin, probed with rabbit MinD polyclonal antibody serum and goat anti-rabbit IgG coupled with horse radish peroxidase (HRP). MinD was visualized using Pierce ECL Western blotting substrate and relative levels were quantified by densitometry using ImageJ (NIH).

## Acknowledgements

We thank Marissa Viola and Ben Piraino for helpful discussions and critical reading of the manuscript, and Janet Atoyan for microscopy and sequencing assistance. Research reported in this publication was supported in part by the National Institute of General Medical Sciences of the National Institutes of Health under Award Number R01GM118927 to J. Camberg. This material is based upon work conducted at a Rhode Island NSF EPSCoR research facility, the Genomics and Sequencing Center, supported in part by the National Science Foundation EPSCoR Cooperative Agreement #OIA-1655221.

## Author contributions

C.J.L., C.E.T. and C.N.F. designed and performed the experiments. C.J.L. and J.L.C. conceptualized the study and analyzed the results. J.C. designed MinD labeling assays, E.C.D. assisted with strain construction and J.J.M. performed electron microscopy. C.J.L. and J.L.C. wrote the manuscript. J.L.C. obtained the funding for the study, supervised the study and managed the project. All authors reviewed, edited and approved the manuscript.

## Additional information

**Supporting information** is available for this paper. (Figure S1, S2, S3, S4, S5, S6)

## References

1. Egan, A. J., and Vollmer, W. (2013) The physiology of bacterial cell division. Annals of the New York Academy of Sciences 1277, 8–28

2. Lutkenhaus, J., Pichoff, S., and Du, S. (2012) Bacterial cytokinesis: From Z ring to divisome. Cytoskeleton 69, 778–790

3. Stricker, J., Maddox, P., Salmon, E. D., and Erickson, H. P. (2002) Rapid assembly dynamics of the Escherichia coli FtsZ-ring demonstrated by fluorescence recovery after photobleaching. Proc. Natl. Acad. Sci. USA 99, 3171–3175

4. Viola, M. G., LaBreck, C. J., Conti, J., and Camberg, J. L. (2017) Proteolysis-Dependent Remodeling of the Tubulin Homolog FtsZ at the Division Septum in Escherichia coli. PloS one 12, e0170505

5. Lutkenhaus, J. (2007) Assembly dynamics of the bacterial MinCDE system and spatial regulation of the Z ring. Annual review of biochemistry 76, 539–562

6. Hu, Z., and Lutkenhaus, J. (1999) Topological regulation of cell division in Escherichia coli involves rapid pole to pole oscillation of the division inhibitor MinC under the control of MinD and MinE. Molecular microbiology 34, 82–90

7. LaBreck, C. J., Conti, J., Viola, M. G., and Camberg, J. L. (2019) MinC N- and C-Domain Interactions Modulate FtsZ Assembly, Division Site Selection, and MinD-Dependent Oscillation in Escherichia coli. Journal of bacteriology 201

8. Lowe, J., and Amos, L. A. (2009) Evolution of cytomotive filaments: the cytoskeleton from prokaryotes to eukaryotes. The international journal of biochemistry & cell biology 41, 323–329

9. Michie, K. A., and Lowe, J. (2006) Dynamic filaments of the bacterial cytoskeleton. Annual review of biochemistry 75, 467–492

10. Hu, Z., and Lutkenhaus, J. (2001) Topological regulation of cell division in E. coli. spatiotemporal oscillation of MinD requires stimulation of its ATPase by MinE and phospholipid. Molecular cell 7, 1337–1343

11. Park, K. T., Wu, W., Battaile, K. P., Lovell, S., Holyoak, T., and Lutkenhaus, J. (2011) The Min oscillator uses MinD-dependent conformational changes in MinE to spatially regulate cytokinesis. Cell 146, 396–407

12. Ghosal, D., Trambaiolo, D., Amos, L. A., and Lowe, J. (2014) MinCD cell division proteins form alternating copolymeric cytomotive filaments. Nature communications 5, 5341

13. Conti, J., Viola, M. G., and Camberg, J. L. (2015) The bacterial cell division regulators MinD and MinC form polymers in the presence of nucleotide. FEBS letters 589, 201–206

14. Huang, H., Wang, P., Bian, L., Osawa, M., Erickson, H. P., and Chen, Y. (2018) The cell division protein MinD from Pseudomonas aeruginosa dominates the assembly of the MinC-MinD copolymers. The Journal of biological chemistry 293, 7786–7795

15. Park, K. T., Du, S., and Lutkenhaus, J. (2015) MinC/MinD copolymers are not required for Min function. Molecular microbiology

16. Camberg, J. L., and Wickner, S. (2012) Regulated proteolysis as a force to control the cell cycle. Structure 20, 1128–1130

17. Camberg, J. L., Viola, M. G., Rea, L., Hoskins, J. R., and Wickner, S. (2014) Location of dual sites in E. coli FtsZ important for degradation by ClpXP; one at the C-terminus and one in the disordered linker. PloS one 9, e94964

18. Camberg, J. L., Hoskins, J. R., and Wickner, S. (2009) ClpXP protease degrades the cytoskeletal protein, FtsZ, and modulates FtsZ polymer dynamics. Proceedings of the National Academy of Sciences of the United States of America 106, 10614–10619

19. Buczek, M. S., Cardenas Arevalo, A. L., and Janakiraman, A. (2016) ClpXP and ClpAP control the Escherichia coli division protein ZapC by proteolysis. Microbiology 162, 909–920

20. Neher, S. B., Villen, J., Oakes, E. C., Bakalarski, C. E., Sauer, R. T., Gygi, S. P., and Baker, T. A. (2006) Proteomic profiling of ClpXP substrates after DNA damage reveals extensive instability within SOS regulon. Molecular cell 22, 193–204

21. Flynn, J. M., Neher, S. B., Kim, Y. I., Sauer, R. T., and Baker, T. A. (2003) Proteomic discovery of cellular substrates of the ClpXP protease reveals five classes of ClpX-recognition signals. Mol. Cell 11, 671–683.

22. Sauer, R. T., Bolon, D. N., Burton, B. M., Burton, R. E., Flynn, J. M., Grant, R. A., Hersch, G. L., Joshi, S. A., Kenniston, J. A., Levchenko, I., Neher, S. B., Oakes, E. S., Siddiqui, S. M., Wah, D. A., and Baker, T. A. (2004) Sculpting the proteome with AAA(+) proteases and disassembly machines. Cell 119, 9–18.

23. Thibault, G., and Houry, W. A. (2012) Role of the N-terminal domain of the chaperone ClpX in the recognition and degradation of lambda phage protein O. J Phys Chem B 116, 6717–6724

24. Thibault, G., Yudin, J., Wong, P., Tsitrin, V., Sprangers, R., Zhao, R., and Houry, W. A. (2006) Specificity in substrate and cofactor recognition by the N-terminal domain of the chaperone ClpX. Proceedings of the National Academy of Sciences of the United States of America 103, 17724–17729

25. Wojtyra, U. A., Thibault, G., Tuite, A., and Houry, W. A. (2003) The N-terminal zinc binding domain of ClpX is a dimerization domain that modulates the chaperone function. J. Biol. Chem. 278, 48981–48990

26. Grimaud, R., Kessel, M., Beuron, F., Steven, A. C., and Maurizi, M. R. (1998) Enzymatic and structural similarities between the Escherichia coli ATP-dependent proteases, ClpXP and ClpAP. J. Biol. Chem. 273, 12476–12481.

27. Wang, J., Hartling, J. A., and Flanagan, J. M. (1997) The structure of ClpP at 2.3 A resolution suggests a model for ATP-dependent proteolysis. Cell 91, 447–456.

28. Camberg, J. L., Hoskins, J. R., and Wickner, S. (2011) The interplay of ClpXP with the cell division machinery in *Escherichia coli*. J. Bacteriol. 193, 1911–1918

29. Hu, Z., Gogol, E. P., and Lutkenhaus, J. (2002) Dynamic assembly of MinD on phospholipid vesicles regulated by ATP and MinE. Proceedings of the National Academy of Sciences of the United States of America 99, 6761–6766

30. Lackner, L. L., Raskin, D. M., and de Boer, P. A. (2003) ATP-dependent interactions between Escherichia coli Min proteins and the phospholipid membrane in vitro. Journal of bacteriology 185, 735–749

31. Singh, S. K., Rozycki, J., Ortega, J., Ishikawa, T., Lo, J., Steven, A. C., and Maurizi, M. R. (2001) Functional domains of the ClpA and ClpX molecular chaperones identified by limited proteolysis and deletion analysis. J. Biol. Chem. 276, 29420–29429

32. Dougan, D. A., Weber-Ban, E., and Bukau, B. (2003) Targeted delivery of an ssrA-tagged substrate by the adaptor protein SspB to its cognate AAA+ protein ClpX. Mol. Cell 12, 373–380

33. Baker, T. A., and Sauer, R. T. (2012) ClpXP, an ATP-powered unfolding and protein-degradation machine. Biochim Biophys Acta 1823, 15–28

34. LaBreck, C. J., May, S., Viola, M. G., Conti, J., and Camberg, J. L. (2017) The Protein Chaperone ClpX Targets Native and Non-native Aggregated Substrates for Remodeling, Disassembly, and Degradation with ClpP. Frontiers in molecular biosciences 4, 26

35. Wawrzynow, A., Wojtkowiak, D., Marszalek, J., Banecki, B., Jonsen, M., Graves, B., Georgopoulos, C., and Zylicz, M. (1995) The ClpX heat-shock protein of Escherichia coli, the ATP-dependent substrate specificity component of the ClpP-ClpX protease, is a novel molecular chaperone. EMBO J. 14, 1867–1877.

36. Levchenko, I., Luo, L., and Baker, T. A. (1995) Disassembly of the Mu transposase tetramer by the ClpX chaperone. Genes Dev. 9, 2399–2408.

37. Barkow, S. R., Levchenko, I., Baker, T. A., and Sauer, R. T. (2009) Polypeptide translocation by the AAA+ ClpXP protease machine. Chemistry & biology 16, 605–612

38. Haeusser, D. P., Lee, A. H., Weart, R. B., and Levin, P. A. (2009) ClpX inhibits FtsZ assembly in a manner that does not require its ATP hydrolysis-dependent chaperone activity. Journal of bacteriology 191, 1986–1991

39. Weart, R. B., Nakano, S., Lane, B. E., Zuber, P., and Levin, P. A. (2005) The ClpX chaperone modulates assembly of the tubulin-like protein FtsZ. Mol. Microbiol. 57, 238–249

40. Salinas, S., Carazo-Salas, R. E., Proukakis, C., Schiavo, G., and Warner, T. T. (2007) Spastin and microtubules: Functions in health and disease. J. Neurosci. Res. 85, 2778–2782

41. Raskin, D. M., and de Boer, P. A. (1999) MinDE-dependent pole-to-pole oscillation of division inhibitor MinC in Escherichia coli. Journal of bacteriology 181, 6419–6424

42. Datsenko, K. A., and Wanner, B. L. (2000) One-step inactivation of chromosomal genes in Escherichia coli K-12 using PCR products. Proceedings of the National Academy of Sciences of the United States of America 97, 6640–6645

43. Maurizi, M. R., Thompson, M. W., Singh, S. K., and Kim, S. H. (1994) Endopeptidase Clp: ATP-dependent Clp protease from Escherichia coli. Methods Enzymol. 244, 314–331.

44. Yakhnin, A. V., Vinokurov, L. M., Surin, A. K., and Alakhov, Y. B. (1998) Green fluorescent protein purification by organic extraction. Protein expression and purification 14, 382–386

45. Conti, J., Viola, M. G., and Camberg, J. L. (2018) FtsA reshapes membrane architecture and remodels the Z-ring in Escherichia coli. Molecular microbiology 107, 558–576

46. Blattner, F. R., Plunkett, G., 3rd, Bloch, C. A., Perna, N. T., Burland, V., Riley, M., Collado-Vides, J., Glasner, J. D., Rode, C. K., Mayhew, G. F., Gregor, J., Davis, N. W., Kirkpatrick, H. A., Goeden, M. A., Rose, D. J., Mau, B., and Shao, Y. (1997) The complete genome sequence of Escherichia coli K-12. Science 277, 1453–1462

