## Supporting information for "Disassembly and degradation of MinD oscillator complexes by *Escherichia coli* ClpXP"

Christopher J. LaBreck, et al., BioRxiv

**Supporting Information**

**Fig. S1. ClpXP degradation reactions with MinC and MinE.** The amount of MinC or MinE after 0 h and 3 h was visualized by SDS-PAGE in reactions containing MinC (6 μM), MinE (6 μM), where indicated, ClpX (1.0 μM), ClpP(1.2 μM), ATP (8 mM) and a regenerating system, as described in *Experimental procedures.*

**Fig. S2. MinD degradation is inhibited by a peptide that binds to the ClpX N-domain.** (A) Degradation of Gfp-ssrA (0.5 μM) was measured by monitoring loss of fluorescence with time in reactions containing ClpP (0.5 μM), ATP (5 mM), ClpX (0.5 μM) or ClpX(ΔN61). (B) MinD degradation rate by ClpXP was determined in reactions containing ClpX (0.75 μM) and ClpP (0.9 μM), fluorescent MinD (4 μM), ATP (8 mM), and increasing concentrations of SspB C-terminal peptide 'XB' (0 to 40 μM). Data from at least three replicates are shown as mean ± SEM. Fluorescent peptides were collected by centrifugation and the rate of degradation was calculated as described in *Experimental procedures*.

**Fig. S3. MinCD copolymer stability in disassembly reactions**. MinCD copolymers were stimulated to assemble with ATP (8 mM) and monitored by 90° light scatter in reactions containing MinC (4 μM), MinD (8 μM) and an ATP regenerating system. Where indicated ClpP

(0.9 μM) or an equivalent amount of buffer was added and copolymers were monitored for an additional 30 min.

**Fig. S4. Activity of MinD(R3E) in vitro.** (A) Copolymer formation by MinD and MinD(R3E) with MinC was compared by monitoring 90° light scatter as described in *Experimental procedures*. Reactions contained MinC (4 μM), ATP (4 mM), and MinD (8 μM) or MinD(R3E) (8 μM), where indicated. Light scatter was monitored for 5 min, ATP was added, and monitored for an additional 30 min. Each curve is representative of at least three replicates. (B) The ability of MinD(R3E) to recruit MinC to SUVs was measured and compared to MinD. Reactions contained SUVs (0.25 mg $ml^{-1}$), ATP (4 mM), MinC (4 μM), MinD (4 μM) or MinD(R3E) (4 μM), as described in *Experimental procedures*.

**Fig. S5. MinD degradation in log phase and stationary phase cultures.** (A) Immunoblots of cells extracts showing relative MinD levels monitored with time in cells that were grown to stationary phase in LB and then treated with spectinomycin (200 μg $ml^{-1}$). Cell extracts from wild type MG1655, *clpX* and *clpP* deletion strains were analyzed by immunoblotting for MinD. (B) Relative MinD levels were monitored with time in cells that were grown to log phase ($OD_{600}$ of 0.3 A.U.) in LB and then treated with spectinomycin (200 μg $ml^{-1}$). Cell extracts from wild type MG1655 and *clpX* deletion strains were analyzed by immunoblotting for MinD followed by densitometry.

**Fig. S6. Intracellular oscillation of Gfp-MinD and Gfp-MinC.** (A) Cells expressing Gfp-MinD from the native locus with and without *clpPX* on the chromosome were visualized by fluorescence microscopy to monitor polar oscillation of Gfp-MinD of log phase cultures. Scale bar is 1 μm. (B) Oscillation rates of Gfp-MinD and (C) Gfp-MinC were measured in at least 20 cells expressing ClpXP from the chromosome (*clpPX*$^{+}$*)*, deleted for *clpPX* (Δ*clpPX*::*kan*)*,* or overexpressing ClpXP from a vector (pClpXP) as described in *Experimental procedures* (p-values are as follows, '*' < 0.05, '**' < 0.005, '****' < 0.0001).

Fig. S1

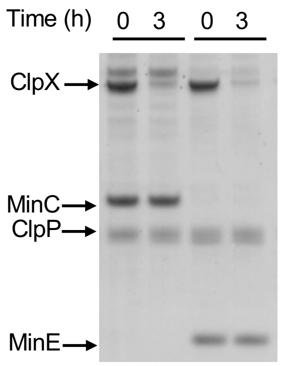

### Fig. S2

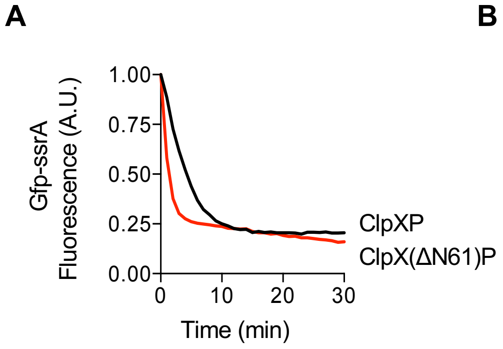
A
Gfp-ssrA Fluorescence (A.U.)
1.00
0.75
0.50
0.25
0.00
0
10
20
30
Time (min)
ClpXP
ClpX(ΔN61)P
B
MinD Degradation Rate (min⁻¹) (normalized to no peptide)
1.00
0.75
0.50
0.25
0.00
0
20
40
Peptide XB (μM)

Fig. S3

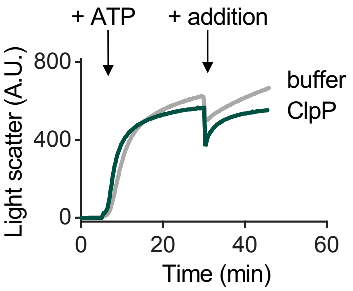
+ ATP
+ addition
buffer
ClpP
Light scatter (A.U.)
800
400
0
0
20
40
60
Time (min)

Fig. S4

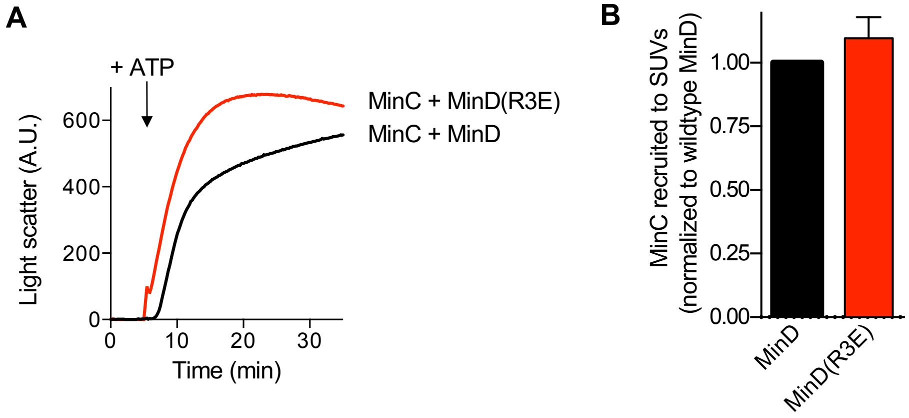
A
+ ATP
Light scatter (A.U.)
600
400
200
0
0
10
20
30
Time (min)
MinC + MinD(R3E)
MinC + MinD
B
MinC recruited to SUVs
(normalized to wildtype MinD)
1.00
0.75
0.50
0.25
0.00
MinD
MinD(R3E)

### Fig. S5

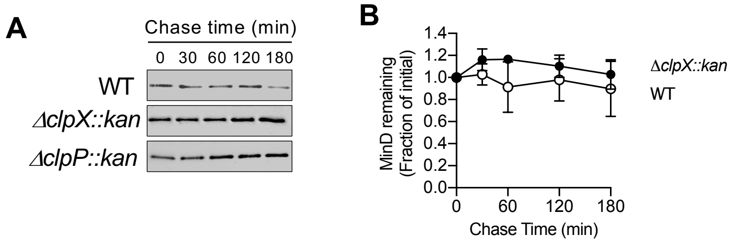
A
Chase time (min)
0
30
60
120
180
WT
ΔclpX::kan
ΔclpP::kan
B
MinD remaining (Fraction of initial)
1.4
1.2
1.0
0.8
0.6
0.4
0.2
0.0
0
60
120
180
Chase Time (min)
ΔclpX::kan
WT

Fig. S6

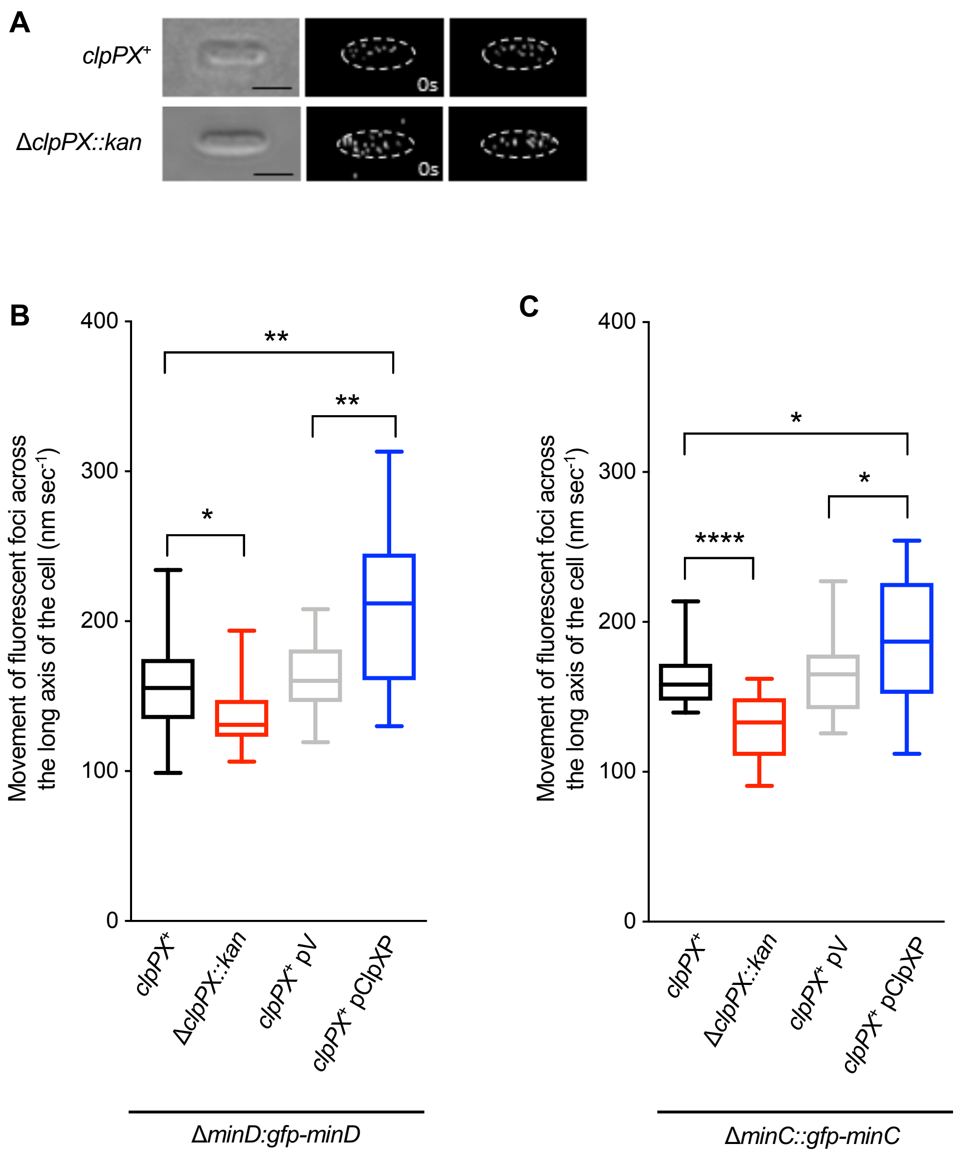
A
clpPX+
ΔclpPX::kan
0s
0s
B
Movement of fluorescent foci across the long axis of the cell (nm sec-1)
400
300
200
100
0
**
**
*
clpPX+
ΔclpPX::kan
clpPX+ pV
clpPX+ pClpXP
ΔminD:gfp-minD
C
Movement of fluorescent foci across the long axis of the cell (nm sec-1)
400
300
200
100
0
*
*
****
clpPX+
ΔclpPX::kan
clpPX+ pV
clpPX+ pClpXP
ΔminC::gfp-minC
